# Metabolomics Ex/GWAS in the Amish reveals novel insights into cardiometabolic disease pathways

**DOI:** 10.1101/2025.08.04.667775

**Authors:** Momodou W. Jallow, Gannie Tzoneva, Bin Ye, Joshua Backman, Valentina A. Zavala, Olukayode Sosina, Manav Kapoor, Niek Verweij, Karl Landheer, Benjamin Geraghty, Joseph Herman, Jonathan Marchini, Adam Locke, Jonas Bovijn, Jonas Nielsen, Luca Lotta, Regeneron Genetics Center, May E. Montasser, Joshua Lewis, Braxton D. Mitchell, Alan R. Shuldiner

## Abstract

We conducted an exome and genome-wide association study (Ex/GWAS) of 1,015 metabolites in serum samples from 5,981 Amish adults. We identified 149 functional or likely functional genetic variants, based on CADD scores significantly associated (*P* < 5.8 x 10^-8^) with 519 metabolite levels, 69 of which were >10-fold enriched in the Amish versus Europeans in gnomAD. We discovered novel associations involving a *PCK2* splice-donor variant (rs138881435) and metabolites important for energy metabolism and mitochondrial function. In UK Biobank participants, this variant was associated with increased longitudinal relaxation time (T1) values on liver MRI suggesting increased inflammation and fibrosis. Furthermore, we found novel associations involving *ENPEP* stop-gain variant (rs33966350) and peptide and amino acid metabolite levels, implicating ENPEP’s roles in blood pressure regulation. Additional known and novel genetic variant-metabolite associations were identified including genes involved in Amish-enriched Mendelian diseases and cardiometabolic traits, providing insights into underlying mechanisms of these disorders.

## BACKGROUND

Metabolomics, the large-scale study of small molecule metabolites within cells, biofluids, tissues, or organisms, provides a direct functional readout of the biochemical and physiological state [1]. Metabolite levels can be influenced by genetic variation, environmental factors, and disease states [2]. Previous metabolomics exome and genome-wide association studies (Ex/GWAS) have identified numerous genetic variant-metabolite associations, linking these findings to human diseases. In the Canadian Longitudinal Survey on Aging cohort, Chen and coworkers identified several gene variant-metabolite and gene variant-metabolite ratio associations, unveiling new biological insights into 12 traits and disease phenotypes including osteoporosis, obesity, inflammatory bowel disease and asthma [3]. A metabolomics GWAS in Finnish men reported 303 novel gene variant-metabolite associations, providing insights into gall stones, cholestasis in pregnancy, and hypertension [4]. Another study conducted in adults from the North Italian Alpine valley identified 85 gene variant-metabolite associations, including 39 novel associations including associations relevant to glutaric acidemia type II and waist circumference [5]. These studies underscore the benefit of using genetically distinct populations in metabolomics Ex/GWAS to facilitate the identification of novel genetic variant associations with metabolites relevant to health and disease.

The Amish population from Lancaster, PA are a genetically distinct closed founder population with a relatively homogeneous lifestyle and well-documented genealogies [6]. As a consequence of a founder effect and genetic drift, they have a higher prevalence of certain genetic variants, many of which have informed health and disease in the Amish and more broadly in other populations [7-9].

In the present study, we conducted untargeted metabolomics on serum samples from Amish adults, with available exome sequence and genome-wide genotype data. Ex/GWAS of 1,015 metabolites in serum from 5,981 individuals, was performed with the aim of identifying novel genetic variants associated with metabolomic traits and exploring their effects on disease and related phenotypes.

## METHODS

### Study cohort

Participants were recruited from the Amish community of Lancaster County, Pennsylvania, USA. The current research entailed a cross-sectional analysis of Amish adults recruited between January 10, 2013 and September 9, 2019 as part of the Amish Wellness Study [10]. Participants provided demographic, diet, medical and medication history, and underwent physical measurements (height, weight, hip and waist circumference, pulse rate and blood pressure). Laboratory measurements of cardiometabolic markers were obtained, including fasting glucose, hemoglobin A1c (HbA1c), cholesterol and creatinine. Serum samples were stored at -80°C for metabolomic profiling. The study was approved by the University of Maryland Baltimore Institutional Review Board, and each participant provided written informed consent prior to enrolment into the study.

### Metabolite measurement, data processing and quality assessments

Untargeted profiling of 1,288 metabolites was conducted on serum samples by Metabolon Inc. (Durham, North Carolina, USA) (**Supplementary Table S1**). Details of the UPLC-MS/MS methods and strict quality control measures were previously described [11].

We evaluated missingness of the non-imputed metabolite data and excluded metabolites and samples with more than 20% missing values (**Supplementary Figure S2**). We did not impute missing values for other metabolites included in the analysis. To reduce skewness, metabolomics data was normalized using rank-inverse normal transformation (RINT) by sex.

To identify potential sources of confounding of metabolite variability, we performed principal component analysis (PCA) on the metabolites using median imputation for missing values. We assessed the impact of age, sex, and body mass index (BMI) on metabolite variability. Additionally, considering that these samples were stored at -80°C for periods ranging from 1 to 11 years (median storage time of 5.86 years) before metabolomics profiling, we examined the effect of sample age and season of sample collection on metabolite variability. These analyses were performed by conducting a linear regression of the covariates against the first two metabolite principal components (PCs) (**Supplementary Figure S1 and Supplementary Table S3**).

### DNA sequencing and genotyping

DNA samples from the 6,270 individuals (3,598 females and 2,672 males) were exome-sequenced and genome-wide genotyped at the Regeneron Genetics Center (RGC) as previously described [7]. Imputation was performed using the Trans-Omics for Precision Medicine program (TOPMed) reference panel which included 1,118 Amish samples. This generated over 9.3 million common variants (MAF ≥ 0.005) and 2.7 million rare variants (MAF < 0.005).

### Exome and genome wide association study (Ex/GWAS) analysis

We performed Ex/GWAS analysis using a linear regression mixed model with REGENIE v3.1.1 [12] on each of the metabolites and samples that passed quality assessments (**Supplementary Figure S2 and Table S2**). Based on examination of the significant determinants of metabolite levels to be included as covariates, we adjusted for age, age^2^, sex, age by sex (age-sex interaction), age^2^ by sex, BMI, season of sample collection, and the first 10 genetic principal components (**Supplementary Tabe S3**). Since many metabolites within the same super pathway exhibited strong correlations (**Figure 1a and Supplementary Table S4**), applying a Bonferroni correction for multiple testing based on the number of metabolites would result in an overly conservative adjustment. Therefore, we opted to use the standard GWAS significance threshold of *P* < 5.0 x 10^-8^ for further evaluation and data mining.

**Figure 1.**
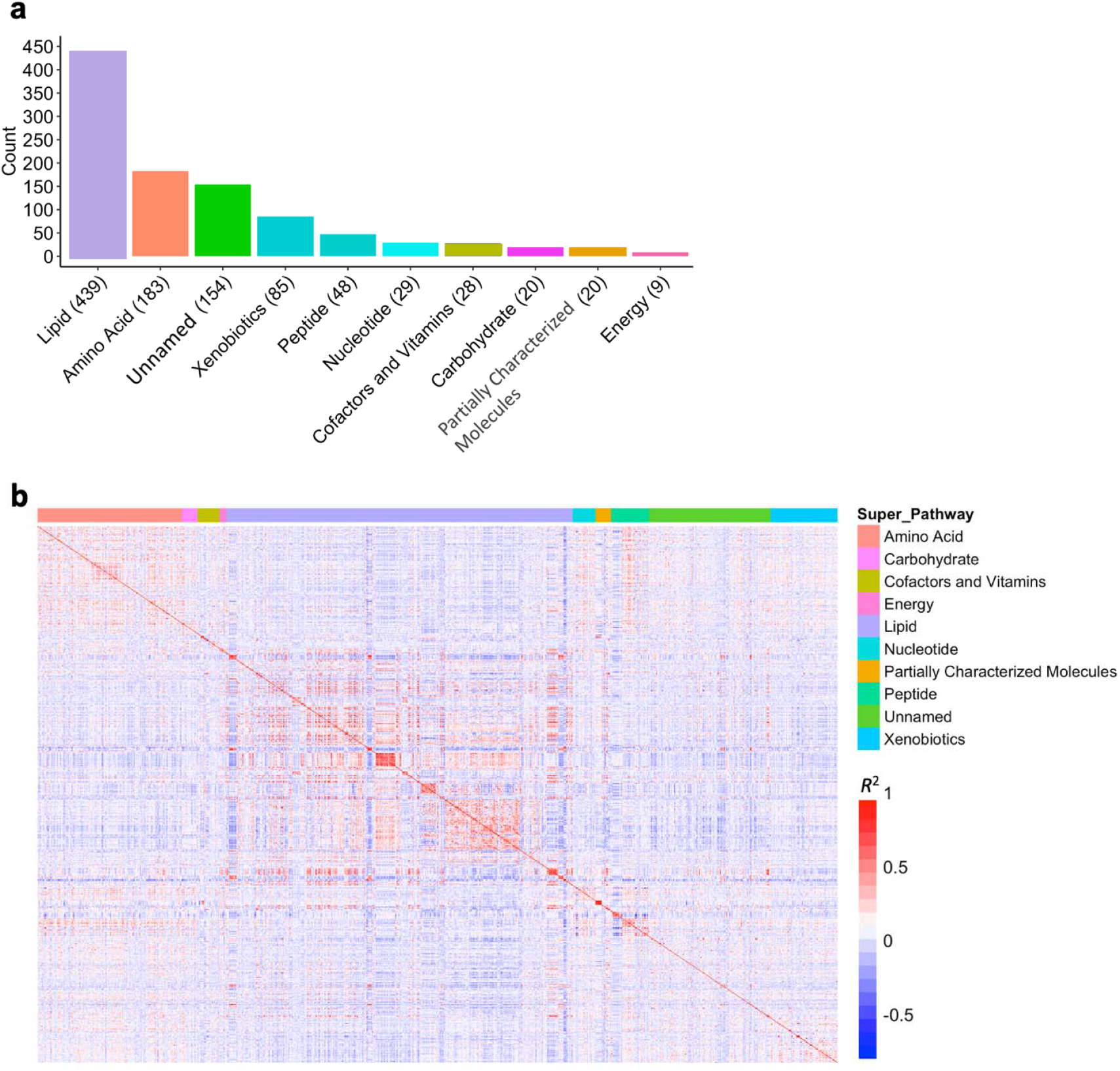
(a) Distribution of the analyzed metabolites across 9 super pathways. (b) Correlation matrix for serum levels of the 1,015 metabolites (see detailed information in **Supplementary Table S4**.)

### Identification of independent association signals with functional or likely functional variants

To identify independent likely causal variants, we examined genetic variant-metabolite association pairs and identified loci with variants that were genome-wide significantly associated with a metabolite and had a CADD (Combined Annotation-Dependent Depletion) score of at least 20. Variants with a CADD score of at least 20 were considered functional or likely functions [13]. We further conducted fine mapping using SuSiE (Sum of Single Effects) to identify genetic variants most likely to causally influence the trait [14]. Regions that were fine mapped were separated by at least 1 Mb. For each locus fitting the above criteria, the genomic region was expanded to cover 1 Mb to ensure comprehensive coverage, and the maximum number of variants within each fine-mapped region was capped at 300,000. These custom parameters were specifically chosen to account for the extended linkage disequilibrium (LD) in this recent (12-14 generation) founder population [15], optimizing the balance between resolution and computational efficiency.

For genomic regions that could not be fine mapped with SuSiE due to extended LD, we implemented a stepwise conditional analysis to identify independent variants within these regions. For each chromosomal region containing at least one genome-wide significant genetic variant-metabolite association (*P* < 5.0 × 10^−8^), the analysis began by conditioning on the variant with the lowest *P*-value. Subsequently, the process iteratively conditioned on the next variant with the lowest *P*-value at the locus. This iterative approach continued until no variant within the region had a *P*-value < 5.0 × 10^−8^.

### Comparing genetic variant-metabolite associations in the Amish with those previously reported

To identify novel associations, we compared our results with published metabolomics Ex/GWAS studies [3, 4, 7, 16-20]. An association was deemed novel if the variant linked to the metabolite had not been previously reported with any phenotype or, although associated with other phenotypes, its specific link to the observed metabolite was not reported. To assess enrichment of the identified genetic variants in our study population, we compared the allele frequencies of these variants in our study population with those observed in non-Finnish Europeans, Finnish, Admixed Americans and African-American populations represented in gnomAD [21].

In addition, we investigated the effects of Amish pathogenic or likely pathogenic (P/LP) variants that are known to cause Mendelian diseases, on metabolite levels. To do this, we queried all the variants-metabolite Ex/GWAS results to identify significant associations involving 228 variants known to be present in the Amish that were considered in ClinVar as P/LP and supported by two stars or higher and an additional 47 variants not in ClinVar identified by the Clinic for Special Children as P/LP (tier 1 genes on the Plain Insight Panel) (Mitchell et al, submitted).

### Follow-up of genetic variant-metabolite associations in the Amish in EHR-linked non-Amish cohorts

We examined the association of metabolites with cardiometabolic traits in the Amish, by regressing the metabolite levels on cardiometabolic traits, adjusting for age, sex, age squared, age-sex interaction (age by sex) and body mass index (BMI). Furthermore, we investigated the impact of the putative functional variants we identified on clinical phenotypes by examining genetic-clinical trait association data from the Amish, in the UK Biobank (UKB) [22].

## RESULTS

Following QC of samples and metabolites, a total of 1,015 metabolites were analyzed (**Figure 1a and Supplementary Table S2**) in 5,981 adult individuals, of whom 57.2% were female (n = 3,419) and 42.8% male (n = 2,562) (**Table 1**).

**Table 1.** Characteristics of Amish participants.

| Baseline variables | All<br>(N = 5981) | Female<br>(n = 3,419) | Male<br>(n = 2,562) |
| --- | --- | --- | --- |
| Age (years)<sup>1</sup> | 40 (18, 93) | 39 (18, 92) | 42 (18, 93) |
| BMI (kg/m<sup>2</sup>)<sup>1</sup> | 25.9 (10.3, 62.0) | 26.1 (10.3, 53.7) | 25.7 (16.6, 62.0) |
| Systolic blood pressure (mmHg)<sup>1</sup> | 112 (78, 236) | 110 (78, 236) | 114 (80, 226) |
| Diastolic blood pressure (mmHg)<sup>1</sup> | 70 (33, 124) | 69 (33.0, 117.0) | 71 (39, 124) |
| Pulse (bpm)<sup>2</sup> | 68.6 (9.9) | 71.2 (10.0) | 65.0 (8.7) |
| Cholesterol (mg/dL)<sup>1</sup> | 202 (64, 514) | 200 (99, 514) | 204 (64, 404) |
| Glucose (mg/dL)<sup>2</sup> | 85.5 (17.0) | 84.5 (17.2) | 87.0 (17.0) |
| Serum sodium (mmol/L)<sup>2</sup> | 138.2 (1.8) | 138.0 (1.9) | 138.5 (1.6) |
| Serum potassium (mmol/L)<sup>2</sup> | 4.2 (0.3) | 4.1 (0.2) | 4.3 (0.3) |

<sup>1</sup> indicates skewed data presented as median and range.
<sup>2</sup> denotes normally distributed data presented as mean and standard deviation.

Levels of many metabolites were highly correlated, especially those belonging to the same pathway (**Figure 1b and Supplementary Table S4**). We found statistically significant effects of age, sex, BMI, and season of sample collection on metabolite levels (all *P* < 0.005), which provided the rationale to include these variables as covariates in genetic analyses (**Supplementary Table S3**). Time of sample storage was not significantly associated with levels of any of the metabolites.

Single gene variant-metabolite associations and fine mapping of likely functional variants From the Ex/GWAS, we identified 550,635 significant (*P*< 5.0 x 10^-8^) single variant-metabolite association pairs for 823 of the 1015 metabolites tested, consisting of 270,435 variants (many variants were associated with more than one metabolite) (**Figure 2**). These variants clustered in 335 genomic regions, defined as genome-wide significant associations located at least 1 Mb apart given extended LD blocks in the Amish beyond what is commonly observed in outbred populations [15]. These significant associations spanned all nine super pathways and unnamed metabolites. Lipid metabolites represent the largest portion, consistent with the higher proportion of lipid metabolites analyzed (**Figure 1a**).

**Figure 2.**
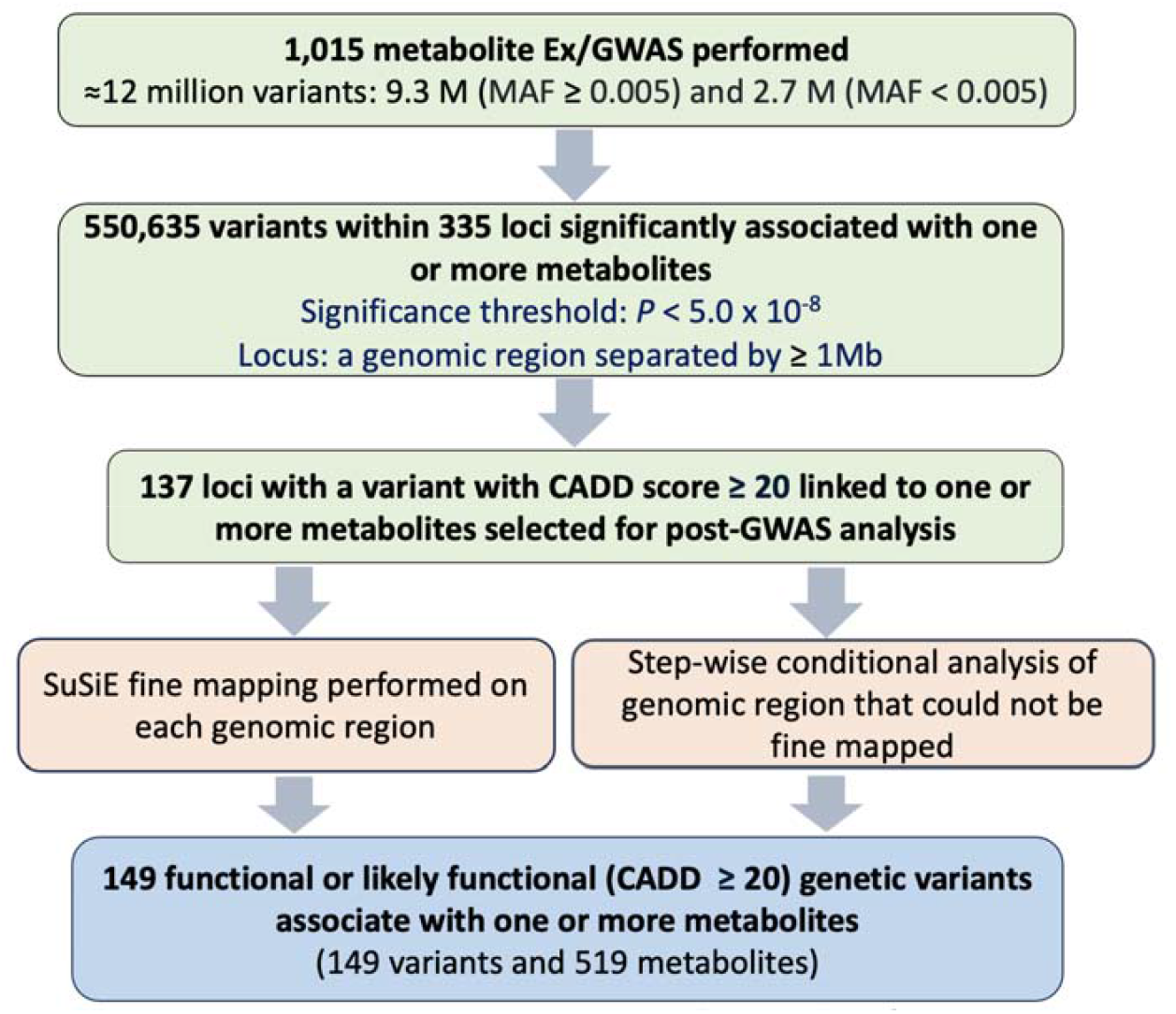
Flow chart illustrating Ex/GWAS analysis and independent functional or likely functional variant identification approach. Post-Ex/GWAS analysis (SuSiE fine-mapping (126 loci) or stepwise conditional analysis (11 loci)) was conducted on loci containing variants that had a CADD score of 20 or higher that were genome-wide significantly associated with one or more metabolites. **CADD =** combined annotation-dependent depletion.

From the 335 genomic loci, we focused post-Ex/GWAS follow-up analysis on 137 loci that contained a genetic variant with a CADD score of at least 20 that was genome-wide significantly associated with one or more metabolites (**Figure 2**). This approach enabled the prioritization of functional or likely functional gene variants driving the observed associations. Fine mapping (126 loci) or stepwise conditional analysis (11 loci which could not be fine mapped likely due to extended Amish LD blocks) identified 149 independent variants with CADD scores of at least 20 significantly associated with one or more metabolites (**Figure 2 and 3a, and Supplementary Tables S5 and S6**).

**Figure 3.**
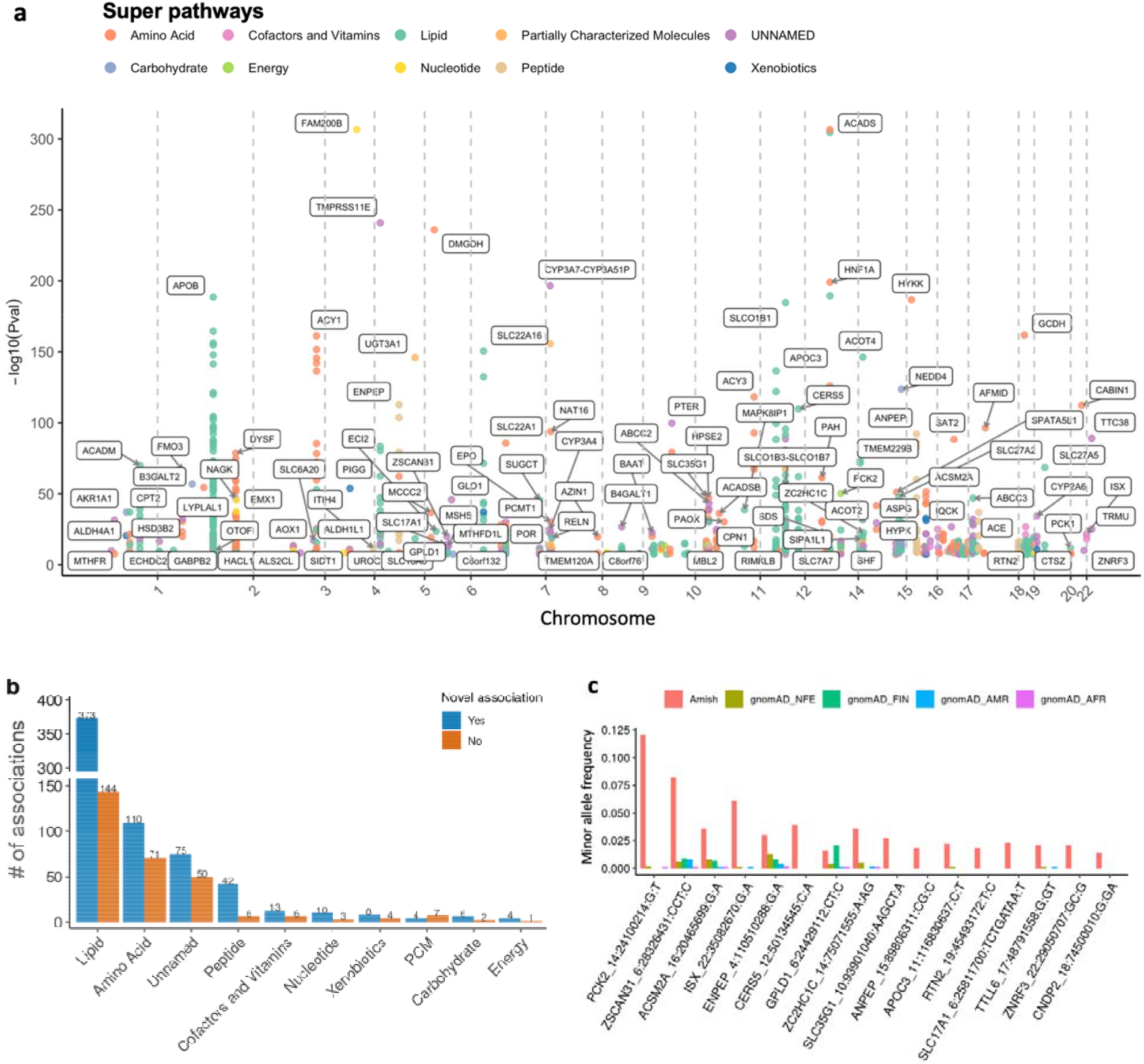
(a) Manhattan plot illustrating 149 functional or likely functional variants that significantly (*P*< 5.0 × 10^-8^) associated with the levels of 519 metabolites. The plot shows statistically significant variant-metabolite associations for each chromosome, while also indicating the super pathway of the associated metabolite. Chromosomes 13, 21 and 23 are absent from the plot because no genome-wide significant associated variants on these chromosomes were found to have a CADD score of 20 or higher. A Manhattan plot showing all the independent variants that associate with one or more metabolites regardless of CADD score is presented in **Supplementary Figure S3**. (b) Comparison of the 939 variant-metabolite association pairs involving 149 functional or likely functional variants, categorized across pathways and stratified by those that have been previously reported vs. those that are new (**Supplementary Table S6**). The X-axis represents the super pathways for each metabolite in the association, while the Y-axis indicates the number of variant-metabolite associations. Blue bars labeled as “Yes” in the legend indicate novel associations, while dark orange bars labeled as “No”, represent previously reported associations. (c) Allele frequency comparisons for 16 pLoF enriched variants significantly associated with one or more metabolites in the Amish vs. four population ancestries in gnomAD [21]. **gnomAD**, Genome Aggregation Database; **NFE**, Non-Finnish Europeans; **FIN**, Finnish; **AFR**, African; **AMR**, Admixed Americans; **PCM**, partially characterized molecules.

### Identification of novel functional or likely functional variant-metabolite associations

From the analysis of 939 genetic variant-metabolite association pairs, we identified 645 novel associations involving 102 putatively functional variants (**Figure 3b and Supplementary Table S7**). Although some of these variants were previously reported to be associated with traits or diseases, their association with these specific metabolites has not been reported previously. Notably, of these 102, 16 are predicted loss-of-function (pLoFs) variants, 13 of which are either only present in the Amish or are >10-fold enriched compared to other populations in gnomAD (**Figure 3c**).

### Amish-enriched genetic variants previously known to be associated with lipid and cardiometabolic traits are associated with known and novel lipid pathway metabolites

We found that the previously reported Amish-enriched *APOB* missense variant on chromosome 2 (rs5742904, p.Arg3527Gln, MAF = 0.067), a cause of familial hypercholesterolemia [23] was linked to 150 metabolites, including cholesterol [Beta (95% CI) = 0.907 (0.840, 0.973), *P* = 1.80×10^-155^], 1-lignoceroyl-GPC (24:0) [Beta (95% CI) = 0.547 (0.481, 0.614), *P* = 6.30×10^-59^] and 1-stearoyl-GPC (18:0) [Beta (95% CI) = 0.423 (0.358, 0.489), *P* = 6.70×10^-37^]. These associations consist of metabolites distributed across five metabolic pathways, with the majority (138/150) related to lipids, and 4 unknown ones (**Supplementary Table S8**).

Also, we found significant genetic variant-metabolite associations involving the previously reported Amish-enriched, cardio-protection-associated *B4GALT1* missense variant (rs551564683, p.Asn352Ser, MAF = 0.061) [8]. This variant was linked to reduced levels of 14 metabolites, including cholesterol [Beta (95% CI) = -0.227 (-0.297, -0.165), *P* = 9.40×10^-12^], and peptide metabolites such as fibrinopeptide A (8-16) [Beta (95% CI) = -0.199 (-0.266, -0.133), *P* = 3.5×10^-9^ and leucylalanine [Beta (95% CI) = -0.221 (-0.291, -0.151), *P* = 6.6×10^-10^] (**Supplementary Table S9**). Notably, 8 out of the *14 B4GALT1* rs551564683-associated metabolites are also significantly associated with APOB rs5742904 in the opposite direction (**Figure 4a**), consistent with opposing roles of these two variants on cholesterol and cardiovascular risk.

**Figure 4.**
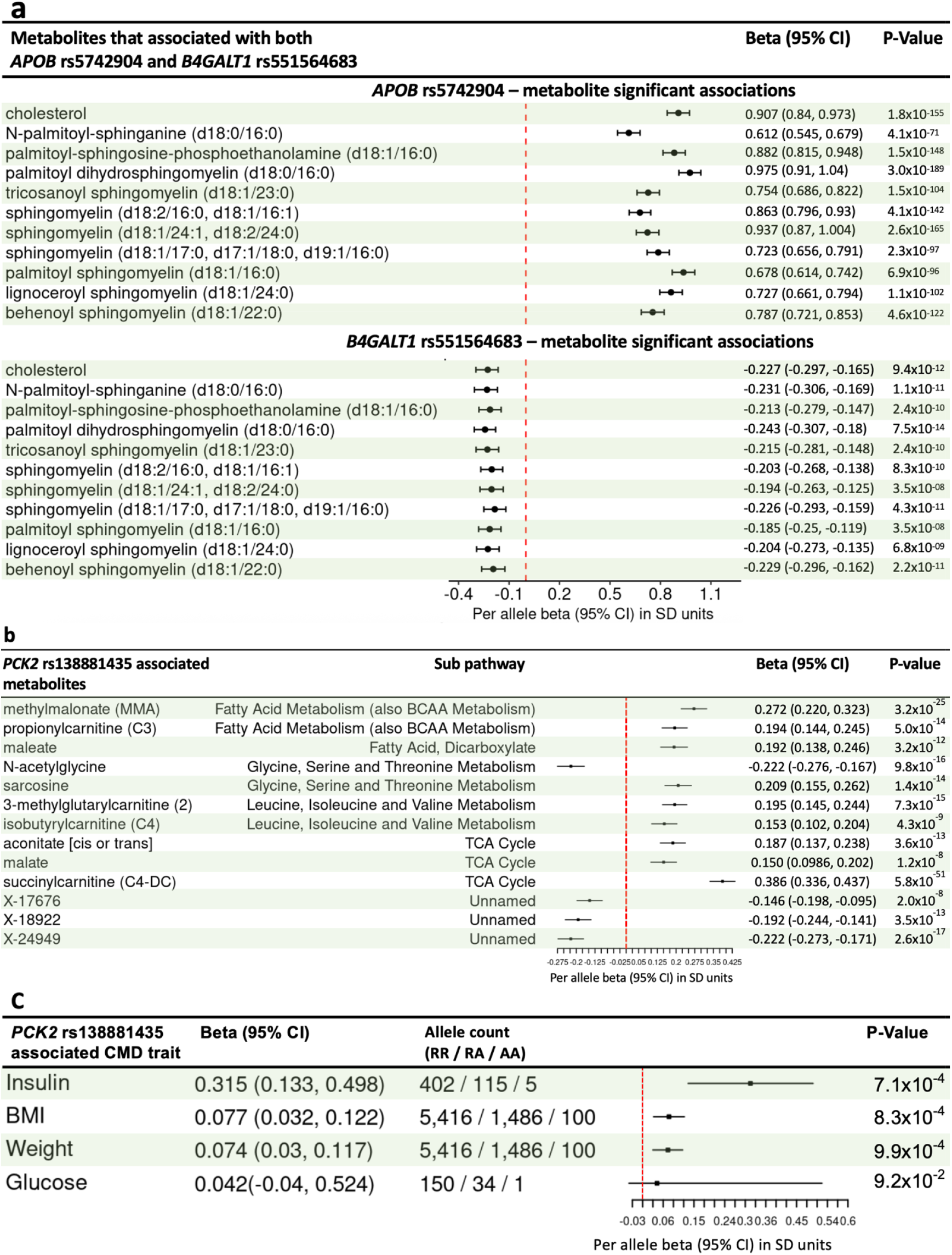
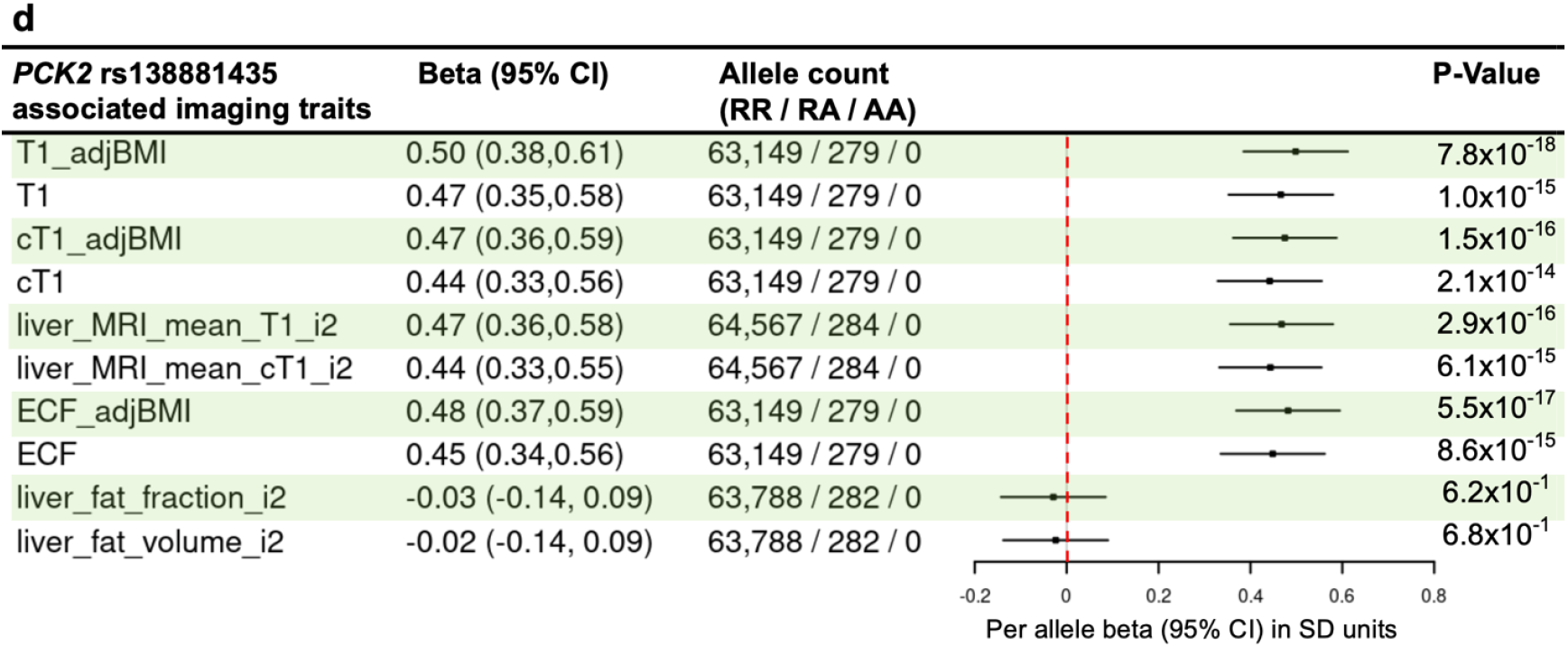
The effects of Amish-enriched gene variants on metabolite levels. (a) Four genetic variant-metabolite associations that overlap between *APOB* rs5742904 (p.Arg3527Gln) and *B4GALT1* rs551564683 (p.Asn352Ser), exhibiting opposing directions of effect. (b) Association between the *PCK2* splice-donor variant rs138881435 (c.1234+1G>T) (AAF = 0.12 with genotype counts RR = 4591, RA = 1266, AA = 86) and the levels of 10 metabolites. (c) Nominally significant association of *PCK2* rs138881435 with fasting insulin, BMI and body weight in the Amish. (d) Association of *PCK2* rs138881435 with liver MRI traits measured in UK Biobank participants. **APOB**, apolopoprotein B; **B4GALT1**, beta-1,4-Galactosyltransferase; **RR**, homozygous for reference allele; **RA**, heterozygous; **AA**, homozygous for alternate allele.; **BMI**, body mass index; **CMD**, cardiometabolic disease; **T1**, longitudinal relaxation time; **AdjBMI**, adjusted BMI; **i2**, second instance; **ECF**, extracellular fluid; **cT1**, corrected T1; **MRI**, magnetic resonance imaging.

The Amish-enriched *APOC3* stop-gain variant (rs76353203, p.Arg19Ter, MAF = 0.022) previously reported to be associated with decreased triglycerides and cardio-protection [24] was associated with the levels of 30 lipid metabolites (**Supplementary Table S6**). Furthermore, we identified significant associations with a previously known Amish-enriched *CERS5* (ceramide synthase 5 gene) splice-donor variant (rs771033566, MAF = 0.039) [8] and the levels of 21 lipid metabolites (**Supplementary Table S6**).

### Amish-enriched pLoF variant in *PCK2* associated with metabolites involved in energy metabolism

We identified a *PCK2* splice-donor variant (rs138881435, c.1234+1G>T) that is highly enriched in the Amish (AAF = 0.12 with 58-fold higher frequency in this population compared to non-Finnish Europeans in gnomAD, 239-fold enrichment compared to Africans, and 379-fold enrichment compared to Admixed Americans in gnomAD (**Supplementary Table S5**). *PCK2* encodes the mitochondrial form of phosphoenolpyruvate carboxykinase (PEPCK-M), which catalyzes the conversion of oxaloacetate to phosphoenolpyruvate [25, 26]. While much is known about the role and regulation of the cytosolic form of PEPCK (PEPCK-C, encoded by *PCK1*) as the rate limiting step in gluconeogenesis, relatively little is known about *PCK2*. Interestingly, we found *PCK2* rs138881435 to be associated with increased levels of several TCA cycle intermediates upstream of oxaloacetate including malate, fumarate, succinylcarnitine (derived from TCA intermediate succinyl-CoA), aconitate and citrate (**Figure 4b**). We also found association with several intermediates of branched chain amino acids and fatty acid metabolism consistent with PEPCK’s known role in glycerolneogenesis (**Figure 4b**). One such intermediate, 3-methylglutarylcarnitine is a recognized biomarker of mitochondrial dysfunction [27], potentially a consequence of perturbation of mitochondrial energy metabolism pathways due to loss-of-function of *PCK2. PCK2* rs138881435 was nominally associated with increased BMI and fasting insulin levels (a marker insulin resistance), but not glucose in the Amish (**Figure 4c**). Although the frequency of *PCK2* rs138881435 is much lower in the UK Biobank, liver MRI data revealed significant association with increased T1 within the liver and increased extracellular fluid (ECF) (**Figure 4d**), reflective of liver tissue alterations, such as inflammation and fibrosis further suggesting metabolic dysfunction [28]. We did not find significant association with liver fat fraction and liver fat volume as measured by MRI (**Figure 4d**).

### The *ENPEP* stop-gain variant rs33966350 (p.Trp413Ter) is associated with peptide and amino acid metabolite levels and hypertension

We identified the previously known *ENPEP* (glutamyl aminopeptidase/aminopeptidase A gene) stop-gain variant (rs33966350, p.Trp413Ter) [29, 30] to be associated with levels of 10 metabolites (8 fibrinopeptides and 2 amino acids) (**Figure 5a and Supplementary Table S6**), which included previously reported associations with six metabolites [16] and four new metabolite associations (**Figure 5b**). Although *ENPEP* rs33966350 (p.Trp413Ter) exists in other populations, its frequency in the Amish (MAF=0.03) is higher compared to other populations (2.2-fold enriched compared to non-Finnish European populations in gnomAD, MAF = 0.012 in EUR) (**Supplementary Table S5**). Fibrinopeptide A (2-15) and fibrinopeptide A, des-ala (1) – both increased in *ENPEP* rs33966350 carriers – were associated with increased systolic and diastolic and mean arterial blood pressure (**Figure 5c**). Interestingly however, these two fibrinopeptides were associated with opposite effects on pulse and ventricular rate, and fibrinopeptide A, des-ala (1) but not fibrinopeptide A (2-15) was associated with lower cholesterol and triglyceride levels. (**Figure 5c**).

**Figure 5.**
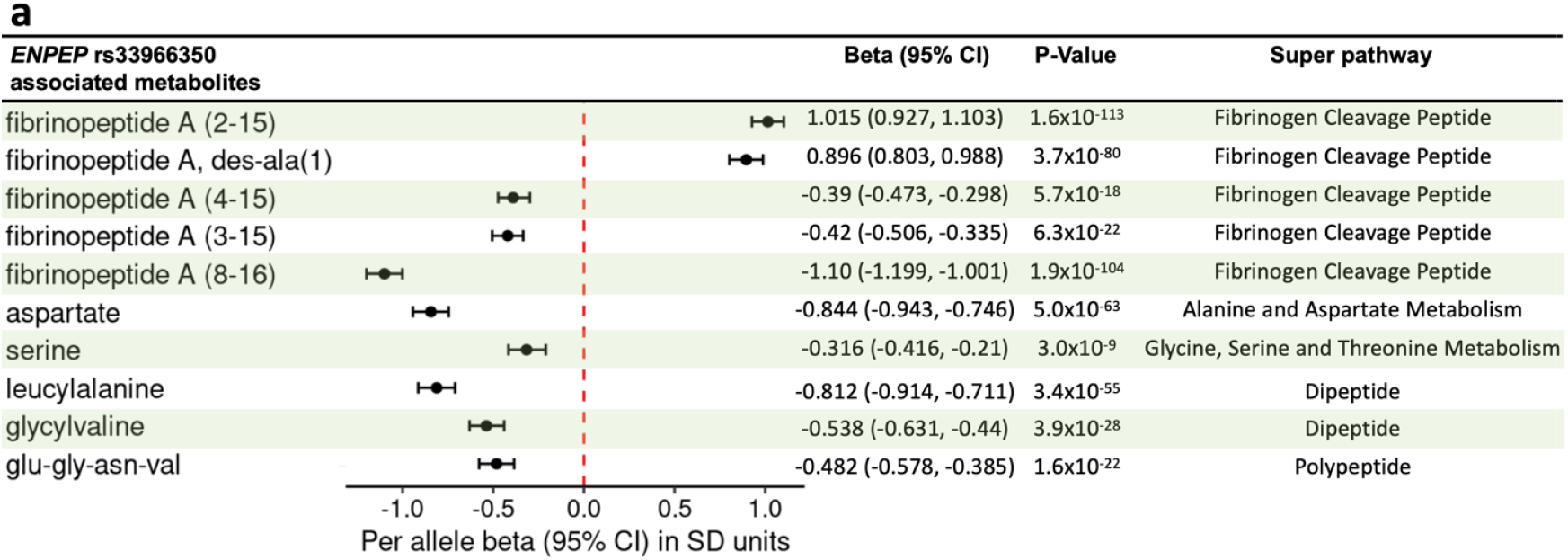
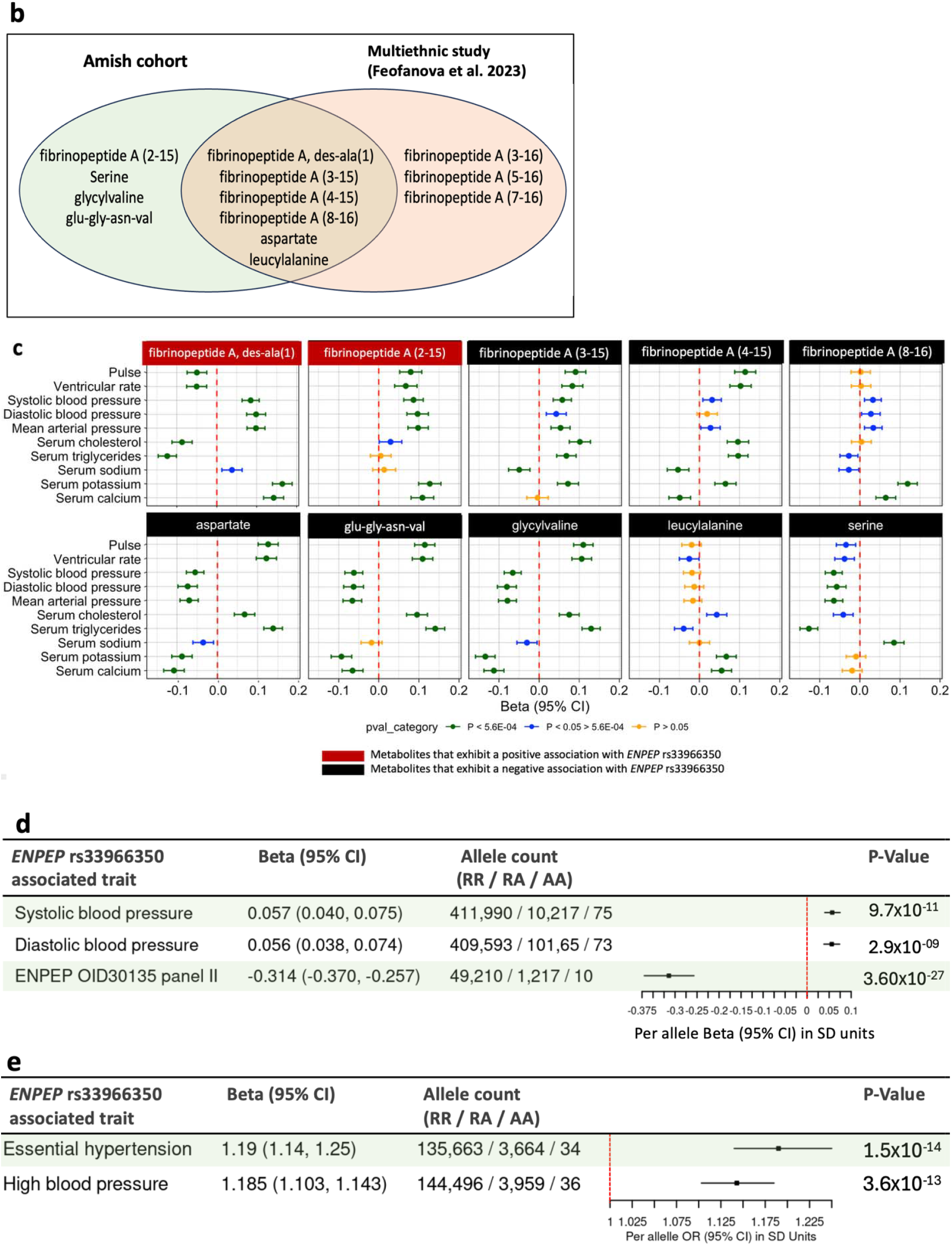
(a) Forest plot showing genome-wide significant associations between *ENPEP* stop-gain variant (rs33966350, AAF = 0.03) and metabolite levels in the Amish cohort. (b) Comparison of *ENPEP* rs33966350-associated metabolites in the present study with those reported by a multiethnic metabolomics GWAS [16]. (c) Association between *ENPEP* rs33966350-associated metabolites and baseline cardiometabolic phenotypes in the Amish cohort. Metabolite names shaded red indicate positive directions of effect of *ENPEP* rs33966350 on the metabolite, while names shaded black indicate negative directions of effect (see **Figure 5a**). Green lines represent statistically significant associations that pass the Bonferroni correction for multiple testing (0.05/100) *P* < 5.0×10, blue lines indicate marginally significant association tests per metabolite below the Bonferroni correction threshold, and yellow lines denote associations with *P* values that are not statistically significant (*P* > 0.05). (d) Associations of *ENPEP* rs33966350 with systolic and diastolic blood pressure in the UK Biobank European cohort [22], and decreased ENPEP protein expression levels in Olink proteomics of approximately 50,000 individuals from the UK Biobank [31]. (e) Associations of *ENPEP* rs33966350 with increased hypertension and high blood pressure in UKB Europeans [22]. **AAF**, alternate allele frequency; **RR**, homozygous for reference allele; **RA**, heterozygous; **AA**, homozygous for alternate allele.

ENPEP is involved in the regulation of systemic arterial blood pressure, which is thought to be through the renin-angiotensin pathway by degrading angiotensin II, which stimulates the secretion of aldosterone, to angiotensin III [32]. To further assess the impact of *ENPEP* stop-gain variant rs33966350 (p.Trp413Ter), we analyzed proteomics data from the UK Biobank (N ≈ 50,000) [31] and found that the variant was significantly associated with decreased ENPEP protein levels (**Figure 5d**). We replicated the previously reported association of ENPEP rs33966350 [30] with increased systolic and diastolic blood pressure (**Figure 5d**) and increased risk of hypertension (**Figure 5e**). We did not find significant association between *ENPEP* rs33966350 and serum or urine potassium or urine potassium-creatinine ratio, serum or urine sodium or sodium-creatinine ratio levels in the UK Biobank (**Supplementary Figure S4**), potentially due to small number of carriers with these phenotypes.

### Known Amish-enriched Mendelian pathogenic or likely pathogenic (P/LP) variants are linked to altered metabolite levels, reflecting the dysregulation of associated genes in both metabolic and non-metabolic diseases

The Amish are enriched for many pathogenic or likely pathogenic (P/LP) variants that cause Mendelian diseases, several of which are metabolic in nature. From the 275 variants investigated, we found association of P/LP mutations in genes causing several metabolic disorders with the expected metabolites (**Supplementary Table S10**). These include for example, a missense mutation (rs121434280, p.Tyr67His) in *ACADM*, known to cause autosomal recessive medium-chain acyl-coenzyme A dehydrogenase deficiency (online Mendelian in man (OMIM) 607008) associated with elevated levels of several fatty acid-containing metabolites; a missense mutation (rs121434367, p.Ala421Val) in *GCDH*, known to cause autosomal recessive glutaric aciduria (OMIM 231670) associated with elevated levels of glutaryl carnitine (C5-DC) and pimeloylcarnitine/3-methyladipoylcarnitine (C7-DC); and a missense mutation (rs119103219, p.Glu99Gln) in *MCCC2*, known to cause autosomal recessive 3-methylcrotonyl-CoA carboxylase 2 deficiency (OMIM 210210) to be associated with elevated levels of isovaline-containing metabolites (**Supplementary Table S10 and S11**). Of note, while these disorders are autosomal recessive, elevations in these metabolites were identifiable in unaffected heterozygous carriers of these mutations.

Other P/LP variants with no clear connection to metabolic disease were found to be associated with metabolite levels that may provide new insights into the pathophysiology of these disorders and/or blood biomarkers for disease risk or therapeutic response in clinical trials (**Supplementary Table S10**). Examples include a stop-gain mutation (rs943680446, p.Arg3400Ter) in *DYNC2H1* associated with asphyxiating thoracic dystrophy 3 (Jeune thoracic dystrophy) (OMIM 613091) associated with decreased levels of several phosphatidylethanolamines; a missense mutation (rs119473030, p.Gly177Ala) in *SLC25A19* linked to Amish microcephaly (OMIM 607196) associated with elevated levels of glutathione and leucine, isoleucine and valine metabolism metabolites; and a frameshift mutation (rs925018791, p.Gly73HisfsTer77) in *SPNS2* known to cause autosomal recessive deafness (OMIM 618457), associated with the decreased levels of palmitoylcarnitine (C16) metabolite (**Supplementary Table S11**).

## DISCUSSION

We performed an exome/genome-wide population-based study to identify genes and functional gene variants associated with metabolite levels and ultimately their effect on human health and disease. The sample consisted of nearly 6,000 Amish individuals from Lancaster County, PA. This population originated from an estimated 400 - 500 founders who immigrated from central Europe to the United States in the mid-18^th^ century and expanded to a population of approximately 43,000 individuals today [33]. While we replicated many genetic variant-metabolite associations previously reported in other European ancestry populations, we focused our analyses on novel associations brought to light due to founder effects, genetic drift and consanguinity. This research offers a comprehensive analysis of metabolomic data from the Lancaster Amish, providing important insights into the genetic architecture of metabolite levels in this population. Indeed, given the substantial fraction of the entire population included in these analyses, we expect to have captured the vast majority of variants and metabolite associations relevant to health and disease in the Lancaster Amish.

In total, we identified 335 genomic loci significantly associated with one or more metabolites. Fine mapping and further analyses identified novel associations of 102 independent genetic variants, 56 of which are enriched more than tenfold in the Amish compared to other populations in gnomAD. Of these 102 variants, 16 are predicted loss-of-function (pLoF), providing further evidence that they are likely causal.

Several enriched genetic variants have been reported in the Amish that influence cardiovascular risk through their effects on lipid homeostasis. These include APOB p.Arg3527Gln which causes familial hypercholesterolemia [23], *B4GALT1* p. Asn352Ser associated with decreased cholesterol (Montasser et al. 2022), and *APOC3* p.Arg19Ter associated with decreased triglycerides (Pollin et al. 2008). Our metabolomics analyses of carriers of these variants provide new mechanistic insights, as well as biomarkers for assessing cardiovascular risk and efficacy of therapeutic and nutritional interventions. For example, *APOB* p.Arg3527Gln was associated with 138 lipid moieties across multiple classes beyond cholesterol and fatty acid metabolism, including increases in ceramides, sphingomyelins, lysophospholipids and endocannabinoids, suggesting that an altered availability of lipid precursors may have more global implications beyond that of cholesterol metabolism and atherosclerotic risk. A recent Mendelian randomization study suggests that genetically predicted *APOB* is associated with decreased healthspan and possibly Alzheimer’s disease [34]. Another study reported ApoB as a novel marker for tau pathology in Alzheimer’s disease [35]. By contrast, in addition to the known association with decreased cholesterol, *B4GALT1* p.Asn352Ser was associated with decreased levels of sphingomyelins, which play diverse roles in membrane function and trafficking and lipoprotein metabolism [36, 37]. *B4GALT1* encodes beta-1,4-galactosyltransferase 1, which is ubiquitously expressed and acts to transfer galactose to specific N-linked glycoprotein substrates [8]. This missense variant has been linked to altered glycosylation patterns of plasma proteins and was also associated with decreased fibrinogen levels [8] likely explaining the observed associations with decreased fibrinopeptides in this study. However, the precise mechanism by which *B4GALT1* p.Asn352Ser influences lipid metabolism is not yet fully understood.

The Amish-enriched triglyceride lowering and cardioprotective *APOC3* p.Arg19Ter variant was associated with decreased levels of phosphatidylethanolamines and increased levels of plasmalogens (**Supplementary Table S6**). Phosphatidylethanolamines play a role in the secretion of lipoproteins in the liver, especially triglyceride-rich very low-density lipoproteins (VLDL) [38]. This observation suggests that in addition to increased conversion of VLDL to LDL observed in stable isotope studies [39], decreased secretion of VLDL by the liver may also contribute to decreased circulating VLDL and cardio-protection observed in p.Arg19Ter *APOC3* carriers. Although the role of increased plasmalogens is less clear, plasmalogens are abundant in heart, brain and myelin sheaths and may play a role in diverse functions including cardiovascular, neurological and immune disorders [40].

Notably, we identified novel associations involving Amish-enriched splice-donor variant (rs138881435, c.1234+1G>T) in the gene encoding the mitochondrial form of PEPCK, *PCK2*. PEPCK converts oxaloacetate, a TCA cycle intermediate to phosphoenolpyruvate, which exits the TCA cycle to participate in other biochemical processes such as gluconeogenesis and glyceroneogenesis [26]. PEPCK also plays important roles in serine biosynthesis and amino acid metabolism [41, 42]. *PCK2* splice-donor variant (rs138881435, c.1234+1G>T) was associated with increased levels of TCA cycle intermediates upstream of its substrate, oxaloacetate suggesting altered energy flux through the TCA cycle. We also found alterations in fatty acid derivatives suggesting derangements in fatty acid and branched chain amino acid metabolism, perhaps through a defect in PEPCK-mediated glyceroneogenesis. Interestingly, *PCK2* rs138881435 was strongly associated with higher levels of 3-methylglutarylcarnitine. This unique acyl-carnitine is derived from 3-methylglutaryl-CoA and cannot be metabolized to yield energy through any known pathway [27]. Increases in 3-methylglutarylcarnitine, as was observed in *PCK2* rs138881435 carriers may be a biochemical marker of mitochondrial dysfunction [27]. *PCK2* rs138881435 associations with these metabolite levels have not been previously reported.

Amish participants in this metabolomics study were generally healthy adults and thus the consequences of *PCK2* rs138881435 on health and disease are unclear, but worthy of further investigation given the pivotal role PEPCK plays in intermediary metabolism. Although this variant is less prevalent in UK Biobank, *PCK2* rs138881435 was strongly associated with increased T1 and extracellular fluid (ECF) on liver magnetic resonance imaging (MRI). This finding indicates increased risk of liver inflammation and fibrosis, possibly due to decreased mitochondrial dysfunction. We did not find any association of this variant with liver fat measures on MRI perhaps reflecting defects in glyceroneogenesis and hepatic triacylglycerol synthesis [28].

Additionally, we found that the hypertension-associated *ENPEP* stop-gain variant (rs33966350, p.Trp413Ter) [29, 30] is associated with levels of peptide and amino acid metabolites. Through the analysis of proteomic data in UK Biobank, we observed that *ENPEP* rs33966350 is associated with decreased levels of ENPEP protein, also known as glutamyl amino peptidase (APA), consistent with a loss-of-function effect. APA is predominantly expressed in the kidneys and small intestines [43]. In the kidney, APA cleaves and inactivates angiotensin II to form angiotensin III. Angiotensin II stimulates the secretion of aldosterone which leads to the retention of sodium and secretion of potassium thus playing a crucial role in fluid and electrolyte balance. Our observed association in UKB Europeans of *ENPEP* rs33966350 with increased blood pressure levels is consistent with increased activity of the renin-angiotensin-aldosterone axis in kidney [44]. In brain, APA generates angiotensin III, which has been shown to have stimulatory effects on systemic blood pressure [45]. Thus, loss of function in APA centrally would be expected to result in decreased angiotensin III and decreased blood pressure. Our observation of increased blood pressure in *ENPEP* rs33966350 carriers suggest peripheral APA activity plays a dominant role in blood pressure regulation relative to its opposing central role.

Given the role of ENPEP in the regulation of blood pressure, modulation of its expression or activity could be targeted to develop new therapies for dysregulation of blood pressure [46] and heart failure [47]. Also, ENPEP has been reported to play a role in vascular inflammation [48], angiogenesis [48], and tumorigenesis and the immune microenvironment [49, 50], potentially offering other opportunities as a therapeutic target for additional indications.

While the *ENPEP* rs33966350-metabolite associations we observed were with small fibrinopeptides, it remains to be determined whether alterations in fibrinogen processing directly influence blood pressure or are simply biomarkers of deranged processing of other peptides such as angiotensin II. Interestingly, two fibrinopeptides that were increased in *ENPEP* rs33966350 carriers had opposite effects on pulse, ventricular rate and cholesterol levels, suggesting that fibrinopeptides may influence these traits.

This work has a number of strengths in identifying enriched genetic variants and metabolite associations that are novel, but we acknowledge several weaknesses. The sample size of approximately 6,000 participants may not have the statistical power to identify associations with rare alleles or alleles of minor or moderate effect size. Participants were adults who were relatively healthy, making difficult our ability to directly connect genetic-metabolite associations to disease traits. Furthermore, we did not have access to medical records but rather more limited, largely cardiometabolic phenotyping obtained for research purposes. The study design was cross-sectional, limiting the ability to discern cause or effect of metabolites on various traits. Future studies may focus on genotype-first call-back studies to explore more deeply the consequences of novel genetic-metabolite associations with biology, health and disease.

In summary, our study demonstrates the power of integrating metabolomics data with genetic analyses in a founder population to uncover novel insights into metabolic regulation. The work underscores the importance of studying diverse populations to capture the full spectrum of genetic variation and its effect on metabolite homeostasis. Future studies should aim to extend these findings in other populations and explore the functional consequences of the identified variants on health and disease.

## Acknowledgements

The Amish Wellness study was approved by the University of Maryland Baltimore Institutional Review Board. Ethical approval for the UK Biobank was previously obtained from the North West Centre for Research Ethics Committee (11/NW/0382). The work described herein was approved by UK Biobank under application number 26041. Informed consent was obtained for all study participants. The authors extend sincere thanks to all the research participants, without whom this research would not be possible. This study is funded by the Regeneron Genetics Center and Regeneron Pharmaceuticals, Inc.

## Supplemental Tables consists of Tables S1-S12 as follows

**Table S1**. List of the 1288 metabolites profiled, their respective super- and sub-pathways, and number and percentage of missing values for each metabolite across all samples assayed.

**Table S2**. List of all the metabolites that passed QC checks and were included in the Ex/GWAS analysis.

**Table S3**. Results of the regression analysis to examine the effects of sample covariates on metabolite PC1 and PC2 derived from metabolite principal component analysis.

**Table S4**. Table S4. Pair-wise correlation of all 1,015 metabolites that passed QC. To facilitate correlation analysis of the metabolites, missing values for each metabolite were imputed using the median of the measured values for that specific metabolite across the entire dataset.

**Table S5**. List of the 149 unique functional or likely functional variants that associate with one or more metabolite levels.

**Table S6**. List of the 939 variant-metabolite associations comprising the 149 independent variants with CADD score of 20 or higher that associate with one or more metabolite levels.

**Table S7**. List of the 649 novel variant-metabolite association pairs featuring 82 independent functional or likely functional variants.

**Table S8**. List of all APOB rs5742904, p.Arg3527Gln (MAF = 0.067) significantly (P < 5.0E-8) associated metabolites (n=150 metabolites), including the super- and sub-pathways of each metabolite.

**Table S9**. A list of all the B4GALT1 rs551564683, p.Asn352Ser, (MAF = 0.061) with 14 metabolites

**Table S10**. List of 37 Amish known pathogenic or likely pathogenic (P/LP) variants from Mitchell et al. (submitted) that associate with the levels of one or more metabolites.

**Table S11**. List of 291 variant-metabolite associations for each of the known Amish P/LP gene variants in Supplementary Table S10.

**Table S12**. List of 2729 associations involving independent variants identified through fine mapping or stepwise conditional analysis (N = 1457 variants) regardless of CADD score, that associate with one or more metabolite levels. This list includes the associations involving the 149 variants with CADD score of 20 or higher in Table S6.

## REFERENCES

1. Wishart, D.S., Metabolomics for Investigating Physiological and Pathophysiological Processes. Physiol Rev, 2019. 99(4): p. 1819–1875.

2. Beger, R.D., et al., Metabolomics enables precision medicine: “A White Paper, Community Perspective”. Metabolomics, 2016. 12(10): p. 149.

3. Chen, Y., et al., Genomic atlas of the plasma metabolome prioritizes metabolites implicated in human diseases. Nat Genet, 2023. 55(1): p. 44–53.

4. Yin, X., et al., Genome-wide association studies of metabolites in Finnish men identify disease-relevant loci. Nat Commun, 2022. 13(1): p. 1644.

5. Konig, E., et al., Whole Exome Sequencing Enhanced Imputation Identifies 85 Metabolite Associations in the Alpine CHRIS Cohort. Metabolites, 2022. 12(7).

6. Khoury, M.J., et al., Inbreeding and prereproductive mortality in the Old Order Amish. II. Genealogic epidemiology of prereproductive mortality. Am J Epidemiol, 1987. 125(3): p. 462–72.

7. Montasser, M.E., et al., An Amish founder population reveals rare-population genetic determinants of the human lipidome. Commun Biol, 2022. 5(1): p. 334.

8. Montasser, M.E., et al., Genetic and functional evidence links a missense variant in B4GALT1 to lower LDL and fibrinogen. Science, 2021. 374(6572): p. 1221–1227.

9. Francomano, C.A., V.A. McKusick, and L.G. Biesecker, Medical genetic studies in the Amish: historical perspective. Am J Med Genet C Semin Med Genet, 2003. 121C(1): p. 1–4.

10. He, S., et al., Prevalence, control, and treatment of diabetes, hypertension, and high cholesterol in the Amish. BMJ Open Diabetes Res Care, 2020. 8(1).

11. Metabolon. Quality Data Results from Quality Sample Handling & Processing. 2025 [cited 2025 05/27]; Available from: https://www.metabolon.com/working-with-us/sample-preparation-handling/.

12. Mbatchou, J., et al., Computationally efficient whole-genome regression for quantitative and binary traits. Nat Genet, 2021. 53(7): p. 1097–1103.

13. Rentzsch, P., et al., CADD: predicting the deleteriousness of variants throughout the human genome. Nucleic Acids Res, 2019. 47(D1): p. D886–D894.

14. Wang, G., et al., A simple new approach to variable selection in regression, with application to genetic fine mapping. J R Stat Soc Series B Stat Methodol, 2020. 82(5): p. 1273–1300.

15. Strauss, K.A. and E.G. Puffenberger, Genetics, medicine, and the Plain people. Annu Rev Genomics Hum Genet, 2009. 10: p. 513–36.

16. Feofanova, E.V., et al., Whole-Genome Sequencing Analysis of Human Metabolome in Multi-Ethnic Populations. Nat Commun, 2023. 14(1): p. 3111.

17. Feofanova, E.V., et al., A Genome-wide Association Study Discovers 46 Loci of the Human Metabolome in the Hispanic Community Health Study/Study of Latinos. Am J Hum Genet, 2020. 107(5): p. 849–863.

18. Ottensmann, L., et al., Genome-wide association analysis of plasma lipidome identifies 495 genetic associations. Nat Commun, 2023. 14(1): p. 6934.

19. Liu, D.J., et al., Exome-wide association study of plasma lipids in >300,000 individuals. Nat Genet, 2017. 49(12): p. 1758–1766.

20. Xu, X., et al., Genetic imputation of kidney transcriptome, proteome and multi-omics illuminates new blood pressure and hypertension targets. Nat Commun, 2024. 15(1): p. 2359.

21. Chen, S., et al., A genomic mutational constraint map using variation in 76,156 human genomes. Nature, 2024. 625(7993): p. 92–100.

22. Bycroft, C., et al., The UK Biobank resource with deep phenotyping and genomic data. Nature, 2018. 562(7726): p. 203–209.

23. Shen, H., et al., Familial defective apolipoprotein B-100 and increased low-density lipoprotein cholesterol and coronary artery calcification in the old order amish. Arch Intern Med, 2010. 170(20): p. 1850–5.

24. Pollin, T.I., et al., A null mutation in human APOC3 confers a favorable plasma lipid profile and apparent cardioprotection. Science, 2008. 322(5908): p. 1702–5.

25. Escos, M., et al., Kinetic and functional properties of human mitochondrial phosphoenolpyruvate carboxykinase. Biochem Biophys Rep, 2016. 7: p. 124–129.

26. Yu, S., et al., Phosphoenolpyruvate carboxykinase in cell metabolism: Roles and mechanisms beyond gluconeogenesis. Mol Metab, 2021. 53: p. 101257.

27. Jennings, E.A., et al., 3-Methylglutarylcarnitine: A biomarker of mitochondrial dysfunction. Clin Chim Acta, 2023. 551: p. 117629.

28. O’Dushlaine, C., et al., Genome-wide association study of liver fat, iron, and extracellular fluid fraction in the UK Biobank. MedRxiv, 2021.

29. Lecluze, E. and G. Lettre, Association Analyses of Predicted Loss-of-Function Variants Prioritized 15 Genes as Blood Pressure Regulators. Can J Cardiol, 2023. 39(12): p. 1888–1897.

30. Surendran, P., et al., Trans-ancestry meta-analyses identify rare and common variants associated with blood pressure and hypertension. Nat Genet, 2016. 48(10): p. 1151–1161.

31. Sun, B.B., et al., Plasma proteomic associations with genetics and health in the UK Biobank. Nature, 2023. 622(7982): p. 329–338.

32. Holmes, R.S., K.D. Spradling-Reeves, and L.A. Cox, Mammalian Glutamyl Aminopeptidase Genes (ENPEP) and Proteins: Comparative Studies of a Major Contributor to Arterial Hypertension. J Data Mining Genomics Proteomics, 2017. 8(2).

33. Lee, W.J., et al., PedHunter 2.0 and its usage to characterize the founder structure of the Old Order Amish of Lancaster County. BMC Med Genet, 2010. 11: p. 68.

34. Martin, L., et al., Mendelian randomization reveals apolipoprotein B shortens healthspan and possibly increases risk for Alzheimer’s disease. Commun Biol, 2024. 7(1): p. 230.

35. Picard, C., et al., Apolipoprotein B is a novel marker for early tau pathology in Alzheimer’s disease. Alzheimers Dement, 2022. 18(5): p. 875–887.

36. Gengatharan, J.M., et al., Altered sphingolipid biosynthetic flux and lipoprotein trafficking contribute to trans-fat-induced atherosclerosis. Cell Metab, 2025. 37(1): p. 274–290 e9.

37. Lee, M., S.Y. Lee, and Y.S. Bae, Functional roles of sphingolipids in immunity and their implication in disease. Exp Mol Med, 2023. 55(6): p. 1110–1130.

38. Vance, J.E., Phosphatidylserine and phosphatidylethanolamine in mammalian cells: two metabolically related aminophospholipids. J Lipid Res, 2008. 49(7): p. 1377–87.

39. Reyes-Soffer, G., et al., Effects of APOC3 Heterozygous Deficiency on Plasma Lipid and Lipoprotein Metabolism. Arterioscler Thromb Vasc Biol, 2019. 39(1): p. 63–72.

40. Paul, S., G.I. Lancaster, and P.J. Meikle, Plasmalogens: A potential therapeutic target for neurodegenerative and cardiometabolic disease. Prog Lipid Res, 2019. 74: p. 186–195.

41. Beale, E.G., B.J. Harvey, and C. Forest, PCK1 and PCK2 as candidate diabetes and obesity genes. Cell Biochem Biophys, 2007. 48(2-3): p. 89–95.

42. Yang, J., S.C. Kalhan, and R.W. Hanson, What is the metabolic role of phosphoenolpyruvate carboxykinase? J Biol Chem, 2009. 284(40): p. 27025–9.

43. Kattula, S., J.R. Byrnes, and A.S. Wolberg, Fibrinogen and Fibrin in Hemostasis and Thrombosis. Arterioscler Thromb Vasc Biol, 2017. 37(3): p. e13–e21.

44. Patel, S., et al., Renin-angiotensin-aldosterone (RAAS): The ubiquitous system for homeostasis and pathologies. Biomed Pharmacother, 2017. 94: p. 317–325.

45. Speth, R.C. and V.T. Karamyan, The significance of brain aminopeptidases in the regulation of the actions of angiotensin peptides in the brain. Heart Fail Rev, 2008. 13(3): p. 299–309.

46. Ferdinand, K.C., et al., Efficacy and Safety of Firibastat, A First-in-Class Brain Aminopeptidase A Inhibitor, in Hypertensive Overweight Patients of Multiple Ethnic Origins. Circulation, 2019. 140(2): p. 138–146.

47. Boitard, S.E., et al., QGC606: A Best-in-Class Orally Active Centrally Acting Aminopeptidase A Inhibitor Prodrug for Treating Heart Failure Following Myocardial Infarction. Can J Cardiol, 2022. 38(6): p. 815–827.

48. Kubota, R., et al., Ischemia-induced angiogenesis is impaired in aminopeptidase A deficient mice via down-regulation of HIF-1alpha. Biochem Biophys Res Commun, 2010. 402(2): p. 396–401.

49. Feliciano, A., et al., miR-125b acts as a tumor suppressor in breast tumorigenesis via its novel direct targets ENPEP, CK2-alpha, CCNJ, and MEGF9. PLoS One, 2013. 8(10): p. e76247.

50. Blanco, L., et al., Altered glutamyl-aminopeptidase activity and expression in renal neoplasms. BMC Cancer, 2014. 14: p. 386.

