## Supplemental Figures S1 to S4 for "Metabolomics Ex/GWAS in the Amish reveals novel insights into cardiometabolic disease pathways"

**Jallow et al.**

**Supplementary notes and figures**

The influence of potential confounding factors on metabolite level variability was investigated through principal component analysis (PCA). PCA was conducted on a dataset comprising 1,015 metabolites, utilizing median imputation to address missing data. This analysis aimed to assess how various factors contribute to the variability in metabolite levels.

We observed that the first two PC explained 14.1% of metabolite variability and 34% of metabolite variability was explained by the first 10 PCs (**Supplementary Figure S1**).

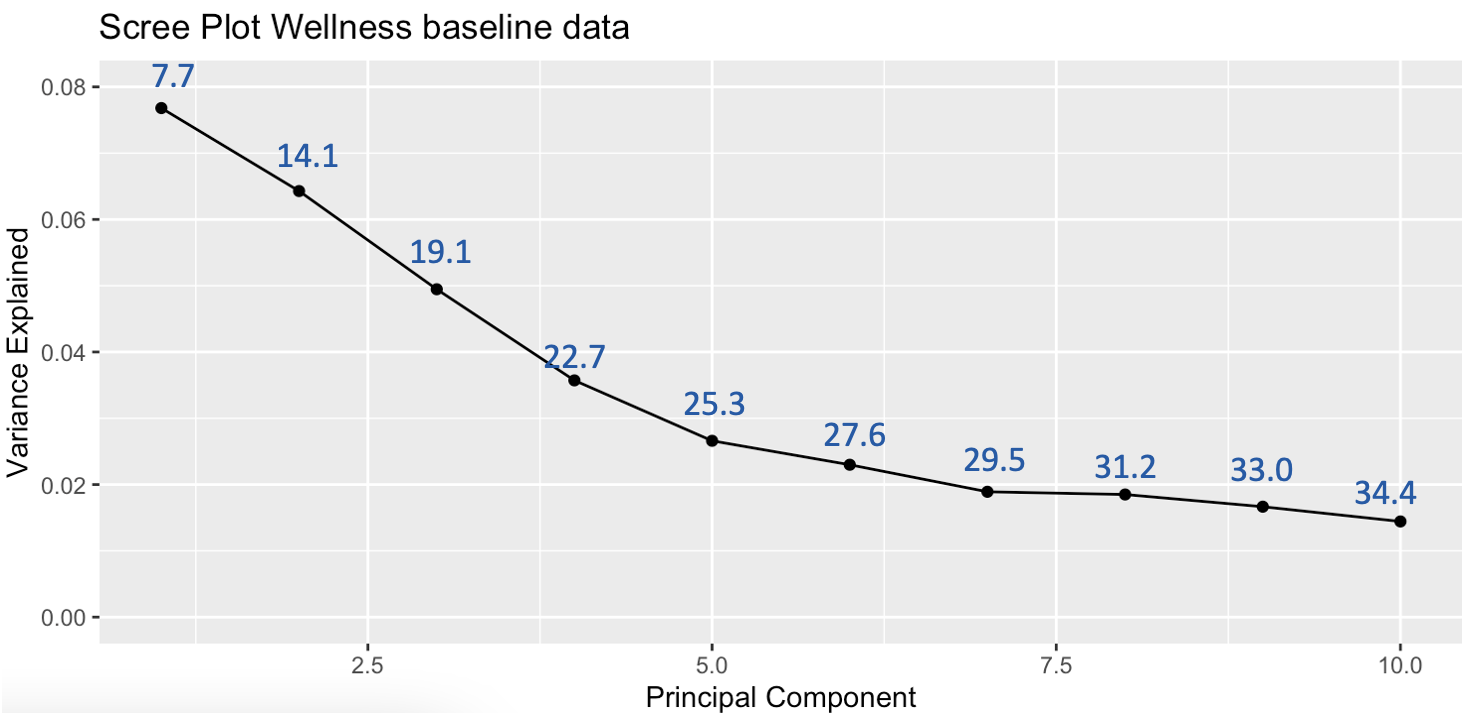

**Supplementary Figure S1.** Plot showing metabolite variability explained by the first 10 metabolite principal components (PCs).

**Evaluation of missingness in the dataset**

Missingness in the metabolomics data ranged from 0 to 99%, with only 329 (25.5%) of the 1,288 metabolites having complete measures (**Supplementary Figure S2a**). Conversely, 90 (7.0%) of the 1,288 metabolites had greater than 90% missing values across all the samples (**Supplementary** **Figure S2a**). Per sample missingness ranged from 9.8% to 28.2% (**Supplementary Figure S2b**), with no sample having complete measure or complete missingness across all metabolites.

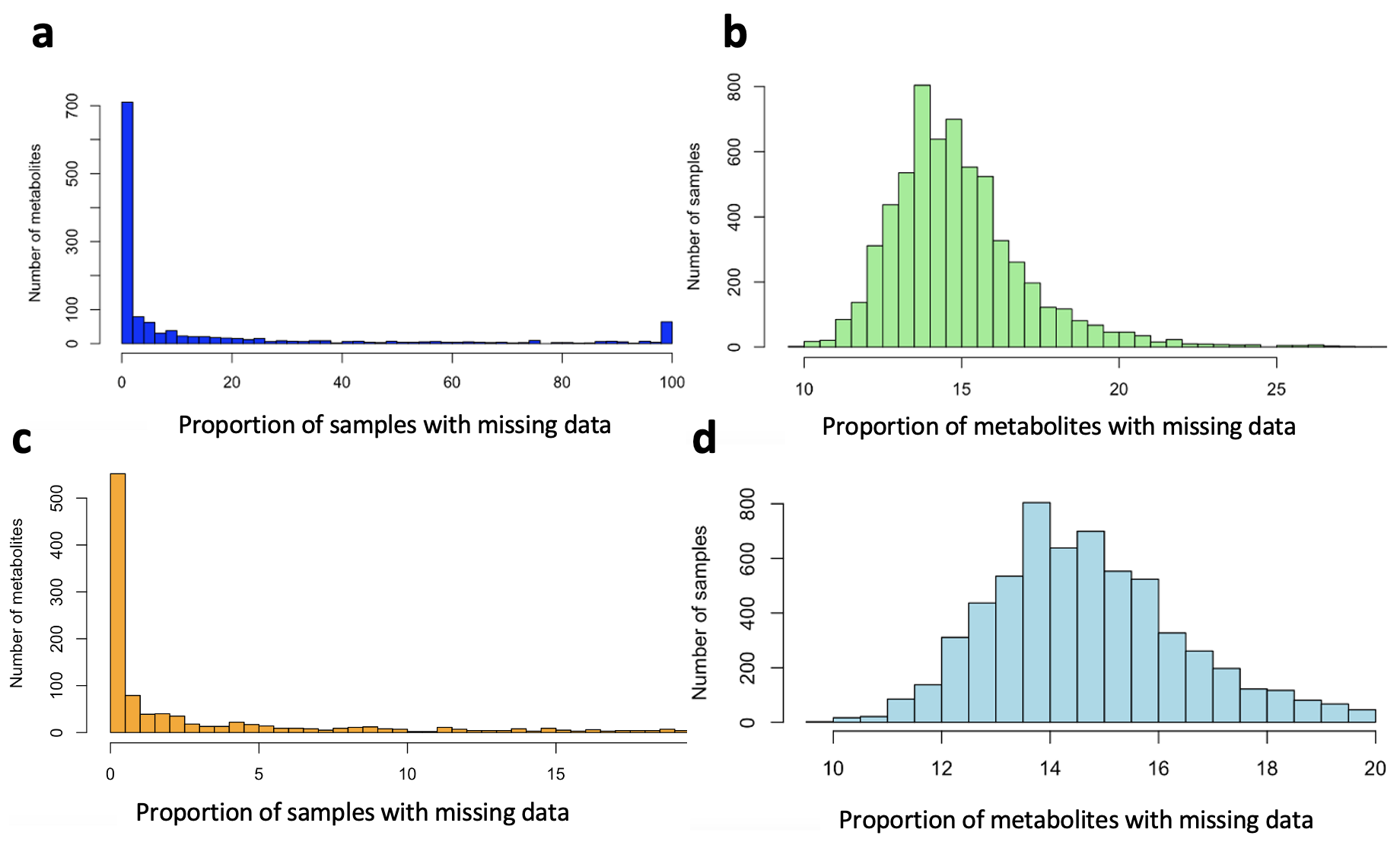

**Supplementary Figure S2**. Distribution of pattern of missingness across metabolites and samples in the study. (**a**) percentage distribution of missingness per metabolite across all samples, (**b**) percentage distribution of sample missingness across all metabolites, (**c**) and (**d**) distribution of metabolites and sample missingness after exclusion of metabolites and samples with greater than 20% missing values, respectively.

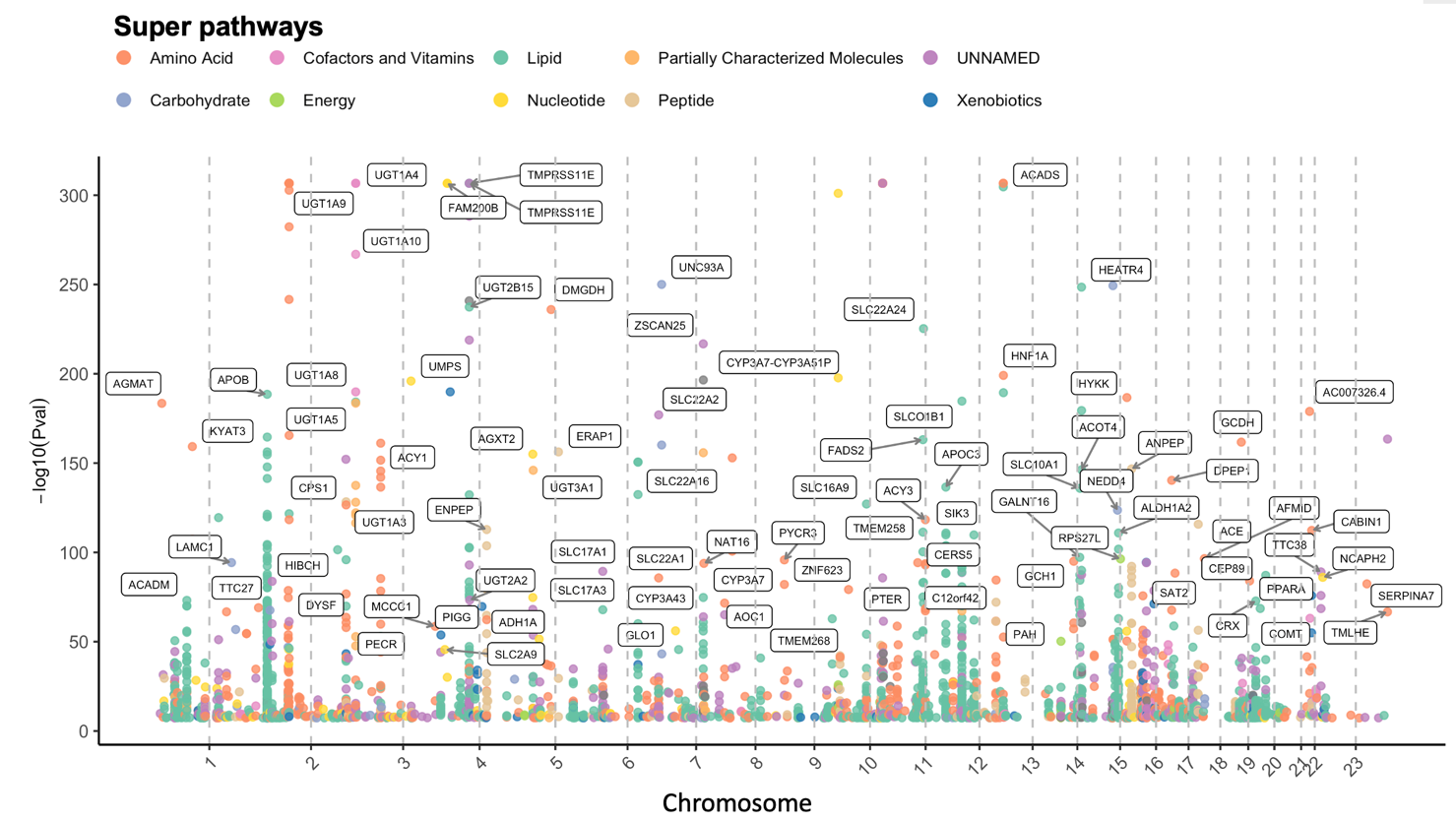

**Supplementary Figure S3.** (a) Manhattan plot illustrating significant independent variant associations across all metabolites regardless of CADD score (1457 genetic variants associate with 664 metabolites). These 1457 genetic variants include the 149 functional or likely functional variants with CADD score of at least 20 presented in **Figure 3a**. Details of the association of these 1457 genetic variants are shown in **Supplementary Table S12**. The plot highlights genome-wide significant genetic variant-metabolite associations on each chromosome and the super pathway to which the associated metabolite belongs.

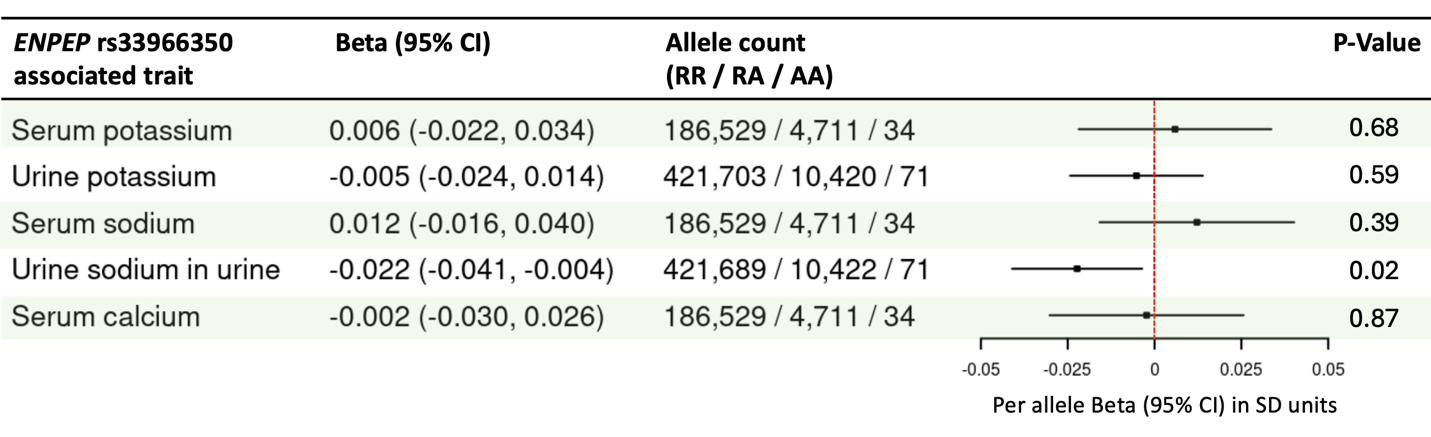

**Supplementary Figure S4.** Lack of association of *ENPEP* rs33966350 with serum or urine potassium, serum or urine sodium and serum calcium in the UK Biobank cohort.
